# Impact of extracellular current flow on action potential propagation in myelinated axons

**DOI:** 10.1101/2024.03.15.585219

**Authors:** Nooshin Abdollahi, Steven A Prescott

## Abstract

Myelinated axons conduct action potentials, or spikes, in a saltatory manner. Inward current caused by a spike occurring at one node of Ranvier spreads axially to the next node, which regenerates the spike when depolarized enough for voltage-gated sodium channels to activate, and so on. The rate at which this process progresses dictates the velocity at which the spike is conducted, and depends on several factors including axial resistivity and axon diameter that directly affect axial current. Here we show through computational simulations in modified double-cable axon models that conduction velocity also depends on extracellular factors whose effects can be explained by their indirect influence on axial current. Specifically, we show that a conventional double-cable model, with its outside layer connected to ground, transmits less axial current than a model whose outside layer is less absorptive. A more resistive barrier exists when an axon is packed tightly between other myelinated fibers, for example. We show that realistically resistive boundary conditions can significantly increase the velocity and energy efficiency of spike propagation, while also protecting against propagation failure. Certain factors like myelin thickness may be less important than typically thought if extracellular conditions are more resistive than normally considered. We also show how realistically resistive boundary conditions affect ephaptic interactions. Overall, these results highlight the unappreciated importance of extracellular conditions for axon function.

**SIGNIFICANCE STATEMENT:** Axons transmit spikes over long distances. Transmission is sped up and made more efficient by myelination, which allows spikes to jump between nodes of Ranvier without activating the intervening (internodal) membrane. Conduction velocity depends on the current transmitted axially from one node to the next. Axial current is known to depend on a variety of features intrinsic to myelinated fibers (e.g. axon diameter, myelin thickness) but we show here, through detailed biophysical simulations, how extracellular conditions (e.g. axon packing density) are also important. The effects ultimately boil down to the variety of paths current can follow, and the amount of current taking alternative paths rather than flowing directly from one node to the next.

## INTRODUCTION

Myelination increases the velocity and efficiency with which axons transmit information. This is because action potentials, or spikes, are regenerated at regularly spaced intervals known as nodes of Ranvier, rather than propagating continuously, as in unmyelinated axons (Frankenhaeuser, 1952; Tasaki, 1952). This so-called saltatory conduction is known to be affected by various morphological and electrophysiological properties intrinsic to the axon. In terms of morphology, conduction velocity increases with axon diameter but depends also on the thickness of the myelin sheath and the distance between nodes (i.e. internode length), both of which co-vary with axon diameter (Brill et al., 1977; Moore et al., 1978). Conduction velocity can also be affected by node length (Arancibia-Cárcamo et al., 2017) and the size of the periaxonal space (Cullen et al., 2021). Sodium channel density at the nodes influences the rapidity of spike regeneration, which in turn also affects conduction velocity (Fields, 2015).

However, “textbook” explanations of saltatory conduction and the role of myelin are not entirely accurate. First, the spike does not occur at one node at a time; instead, repolarization is not yet complete at upstream nodes before that spike re-initiates at downstream nodes, meaning a spike (at slightly different phases of its evolution) exists across several nodes. Second, myelin does not provide tight insulation that altogether prevents the axon membrane in the internode region from charging. Contrary to the “node-to-node” model, which predicts that the passive electrical response of the axon should be fast because the input resistance and capacitance of the nodes are low and capacitance of the myelin sheath is low, Barrett and Barrett (1982) demonstrated a slow depolarizing afterpotential caused by the internodal axon membrane becoming charged during spike propagation, and discharging after the spike has passed. This implies charge separation across the internodal axon membrane, which in turn implies a space under the myelin. Whether current flows laterally across the myelin or longitudinally along the submyelin (periaxonal) space has been debated (Blight & Someya, 1985); these paths are not mutually exclusive. This prompted the development of double-cable axon models (Blight, 1985), the most popular of which is arguably the MRG model (McIntyre et al., 2002). The nuances of saltatory conduction continue to be actively investigated; for example, Cohen et al. (2020) combined structural and functional measurements with sophisticated computational modeling to reconcile several observations. But one assumption of single- and double-cable models is that extracellular space is ultimately connected to ground, resulting in an infinite extracellular conductivity. That boundary condition is probably accurate for isolated fibers but may be inaccurate for fibers packed into an intact nerve. A corollary of this is that transmembrane voltage is often calculated under the assumption that extracellular voltage is zero, but this too is inaccurate under various conditions.

A spike propagating down an axon can be recorded extracellularly because it alters the extracellular potential. This change in extracellular potential also affects nearby axons, causing ephaptic interactions (Jefferys, 1995). Ephaptically mediated voltage changes tend to be very small but can nonetheless advance or delay spike propagation, which can contribute to synchronized firing (Bokil et al., 2001; Han et al., 2018; Reutskiy et al., 2003). But the sensitivity of ephaptic interactions on current flow in the extracellular space, including the importance of factors like packing density, has not been thoroughly investigated. Separate from ephaptic effects, efforts to infer axon parameters by fitting models to physiological measurements may be compromised by misapproximating boundary conditions.

In this study, we examined effects of extracellular electrical properties on saltatory spike propagation by varying boundary conditions in modified double-cable axon models. Using these models, we demonstrate that extracellular conditions influence conduction velocity and ephaptic interactions, as well as the reliability of spike propagation under demyelinating conditions. We also demonstrate how extracellular conditions influence the impact of other factors on conduction velocity.

## METHODS

Simulations were conducted in Neuron 8.0 (Hines & Carnevale, 1997) and Python using NetPyNE (Dura-Bernal et al., 2019). We adapted the “MRG” axon model developed by McIntyre et al. (2002). The MRG model is based on motor axons but has been widely used to a wide range of axons peripherally and centrally (Capllonch-Juan & Sepulveda, 2020; Howell et al., 2015; Jones et al., 2021; McIntyre et al., 2004; Mirzakhalili et al., 2020; Sagalajev et al., 2023).It is a double-cable model that includes two layers of extracellular space with the outermost space (referred to here as the extramyelin layer) attached to ground (condition 1). We modified the original single-fiber model by disconnecting extramyelin compartments from ground and connecting, via transverse resistance, the outermost layer of extracellular space above each node (adjacent to the extramyelin layer above paranodes) to ground to simulate conditions 2 and 3. In multi-fiber models, the outermost layer of extracellular space above each node is connected via transverse resistance to the equivalent layer of an adjacent node on a nearby fiber and to a perineurium, which represents a boundary defining a fascicle of fibers. To model boundary conditions in multi-fiber simulations, boundary cables including resistances were placed near the axons. The transverse resistances between the axons were calculated as follows (Capllonch-Juan & Sepulveda, 2020):

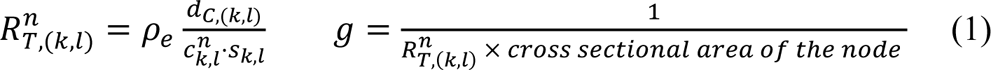

where *ρ*_*e*_ is the resistivity of endoneurium (the interfascicle tissue between the fibers) in unit (Ω ⋅ cm), *d*_*C*,(*k*,*l*)_ is the edge-to-edge distance between axons *k* and *l*. 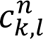 is the length (along the z-axis) of the transverse resistor number n between axons *k* and *l*. *S_k,l_* is the mean of the axon’s diameters. The radius of the fiber used in simulation is 4 microns.

To calculate the transverse resistances between the axons and the boundary cables, *d*_*C*,(*k*,*l*)_ and *ρ*_*e*_ in equation 1 is replaced with the thickness (4.7 ×10^−4^ cm) and the resistivity of perineurium (1.136 ×10^5^ Ω.cm) respectively. Resistivity of endoneurium is 1211 Ω.cm.

Longitudinal resistances are calculated as follows:

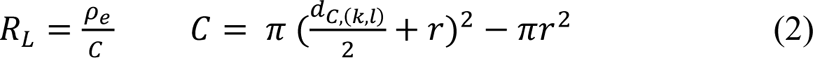

where *C* is the cross-sectional area of the extracellular space around each fiber.

For aligned fibers, only the nodes of Ranvier are transversely connected to each other. For misaligned fibers, the nodes are connected to nearby compartments. To investigate the effects of anatomical properties on ephaptic coupling, the number of fibers was increased to 20. All fibers have the same intrinsic properties and are connected to each other via transverse resistances with the values calculated by equation 1. The longer the distance between the fibers the larger the transverse resistance connecting them. Also, the fibers are all connected to boundary cables with different transverse resistance values. The fibers on the edge have smaller transverse resistance to the boundary cables.

All model code will be made available on ModelDB upon publication.

## RESULTS

### Extracellular properties impact conduction velocity

Myelinated axons are commonly simulated using a double-cable model (McIntyre et al., 2002), which consists of intracellular and extracellular cables along which current can flow. The extracellular space is subdivided into the submyelin (periaxonal) space and the extramyelin space (**Fig. 1A**). One assumption of conventional double-cable models is that the extramyelin space is connected to ground. Under this condition, extracellular current is promptly shunted to ground rather than flowing longitudinally (parallel to the axon) and/or laterally (orthogonal to the axon) depending on the resistance of various pathways. This may or may not accurately reflect what happens for a real fiber depending, for example, on whether a fiber is positioned on the edge or in the middle of a bundle (fascicle) of fibers, and how densely those fibers are packed (see cartoons on right side of Fig. 1B). In particular, the spread of extracellular current may experience much greater resistance than is experienced when shunted straight to ground.

**Figure 1.**
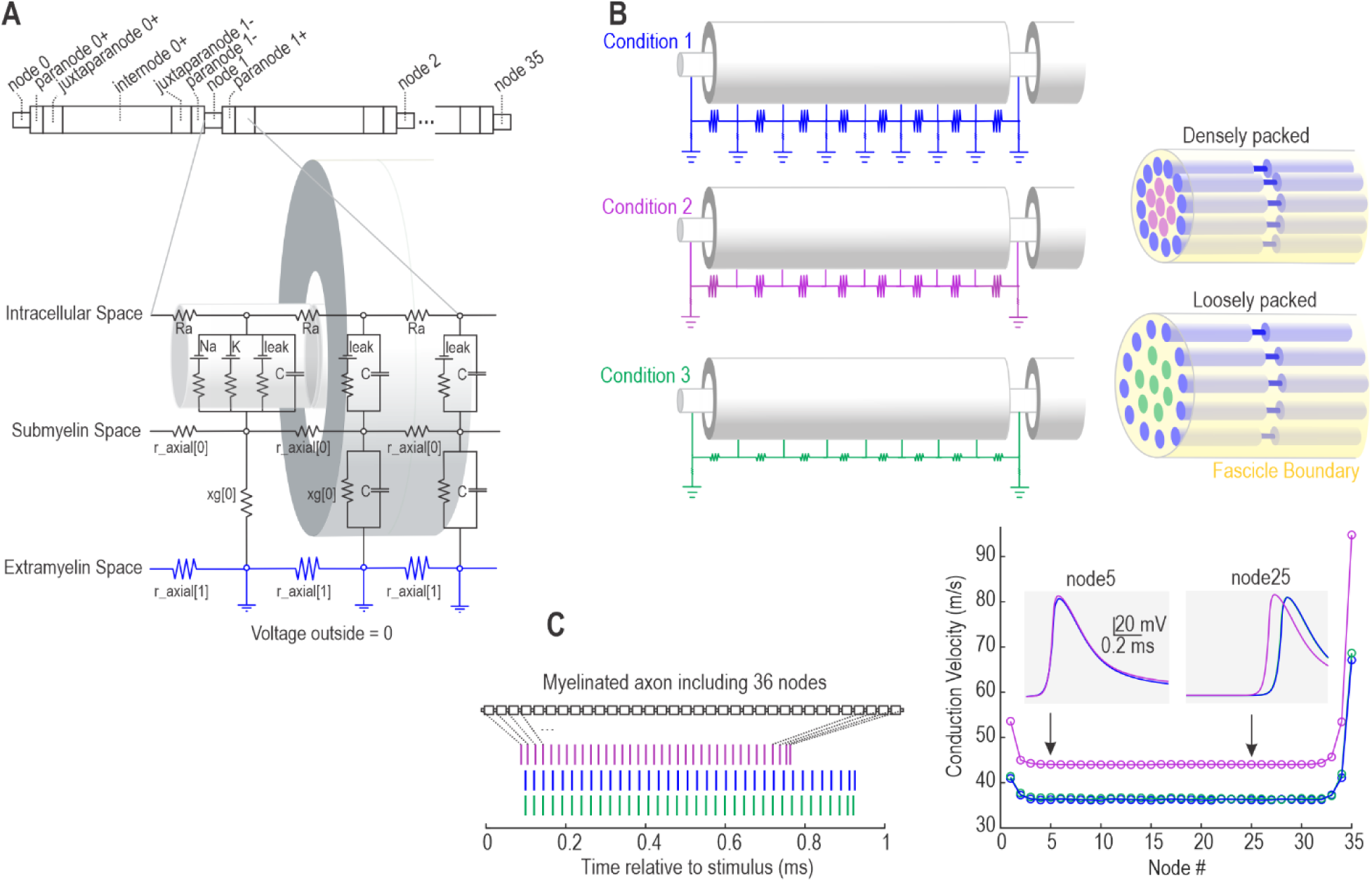
Extracellular properties affect axon conduction velocity. **A.** Equivalent electrical circuit of the conventional double-cable model. The extracellular space is subdivided into submyelin and extramyelin spaces, which are separated by myelin in the internode, juxtaparanode, and paranode regions. The extramyelin space for all regions is connected to ground, thus forcing extracellular voltage to be zero. Cartoon at top shows numbering of nodes from left (0) to right (35); all other sections are numbered relative to nodes, where + and – mean to the right or left, respectively. Section lengths are not drawn to scale. **B**. Three extracellular conditions. Condition 1 (blue) represents the conventional model (see panel A), which approximates a fiber located at the edge of a fascicle with absorptive boundaries. Conditions 2 (purple) and 3 (green) represent a fiber positioned in the middle of a fascicle that is densely packed (Condition 2) or loosely packed (Condition 3); packing density is parameterized by varying the longitudinal resistance of the extramyelin space. Cartoons on the right depict the multi-fiber geometry, using the same color scheme as on the left. **C.** After initiating a spike at the left end of the fiber, the time of spike regeneration at each of the 36 nodes was recorded (left panel). Conditions are colored like in B. Conduction velocity was calculated from the spike time interval between consecutive nodes (right panel). Conduction velocity is higher at each end of the fiber because of longitudinal boundary conditions. Conduction velocity (based on middle nodes) was higher in condition 2 than in conditions 1 and 3. Shaded insets show the spike waveforms at nodes 5 and 25.

To explore this, we modeled a single myelinated fiber in three conditions **(Fig. 1B)**. Condition 1 is a conventional double-cable model insofar as all sections of the extramyelin space are connected to ground; this condition reasonably approximates a fiber located at the edge of a fascicle with a highly absorptive (i.e. conductive, non-resistive) boundary. In Condition 2, the internode sections of the extramyelin space are disconnected from ground to approximate a fiber situated in the middle of a fascicle; only the extracellular space overlying nodes is connected to ground, therein simulating conditions in which the alignment of nodes across adjacent fibers offers a relatively low resistance path to the fascicle edge; misaligned nodes are considered later. In Condition 3, the longitudinal extramyelin resistivity is reduced to model a fiber within a fascicle with low packing density. All intrinsic properties of the fiber used in different conditions are identical, meaning only extracellular factors differ. The conditions simulated here, for a single fiber, are validated later using multi-fiber simulations (see below).

Using the same stimulus to evoke a spike at one end of the fiber (node 0), the spike initiation time was determined for each node, for each condition, and the interval between spikes at consecutive nodes was used to measure conduction velocity **(Fig. 1C)**. Conduction velocity is higher at each end of the fiber (because of longitudinal boundary effects) and we therefore report conduction velocity based on intermediate nodes (i.e. from the flat region of the graph). A fiber in the middle of a densely packed fascicle (Conditions 2) had a higher conduction velocity than a fiber located at the fascicle edge (Condition 1) or in the middle of a loosely packed fascicle (Condition 3). These results clearly suggest that extracellular conditions influence conduction velocity.

### Effects on conduction velocity are explained by changes in extracellular current

To elucidate the biophysical basis for effects reported in Figure 1, we calculated the axial current and node transmembrane current as well as the submyelin, transmyelin and extramyelin currents between nodes 14 and 15, which are located in the middle of the axon. The spike regenerated at node 14 under each extracellular condition is aligned in time so that any difference in spike regeneration at node 15 is due to differences in current flow between nodes 14 and 15, and not because of effects upstream of node 14.

Figure 2A compares current flow between Condition 1 (slow) and Condition 2 (fast). In Condition 2, the membrane of node 15 charges slightly faster, evident by a slightly larger upward deflection in the transmembrane current, followed by earlier activation of Na channels, evident by the earlier downward deflection (Fig. 2B). To account for the faster charging of node 15 in Condition 2 relative to Condition 1, we computed the axial current leaving node 14 and subsequently received by node 15 (Fig. 2C). The results revealed that as a spike propagates, axial current is less attenuated in Condition 2, leading to slightly more axial current reaching node 15 than in Condition 1 The greater attenuation of axial current in Condition 1 is because all sections of the extramyelin space are connected to ground, which encourages current leakage across the myelin (Fig. 2D **left**). Current that leaks across the myelin does not flow longitudinally in the extramyelin space but is, instead, shunted to ground (Fig. 2D **middle)**. Since less current leaks across the myelin in Condition 2, more current flows along the submyelin space (Fig. 2D **right**).

**Figure 2.**
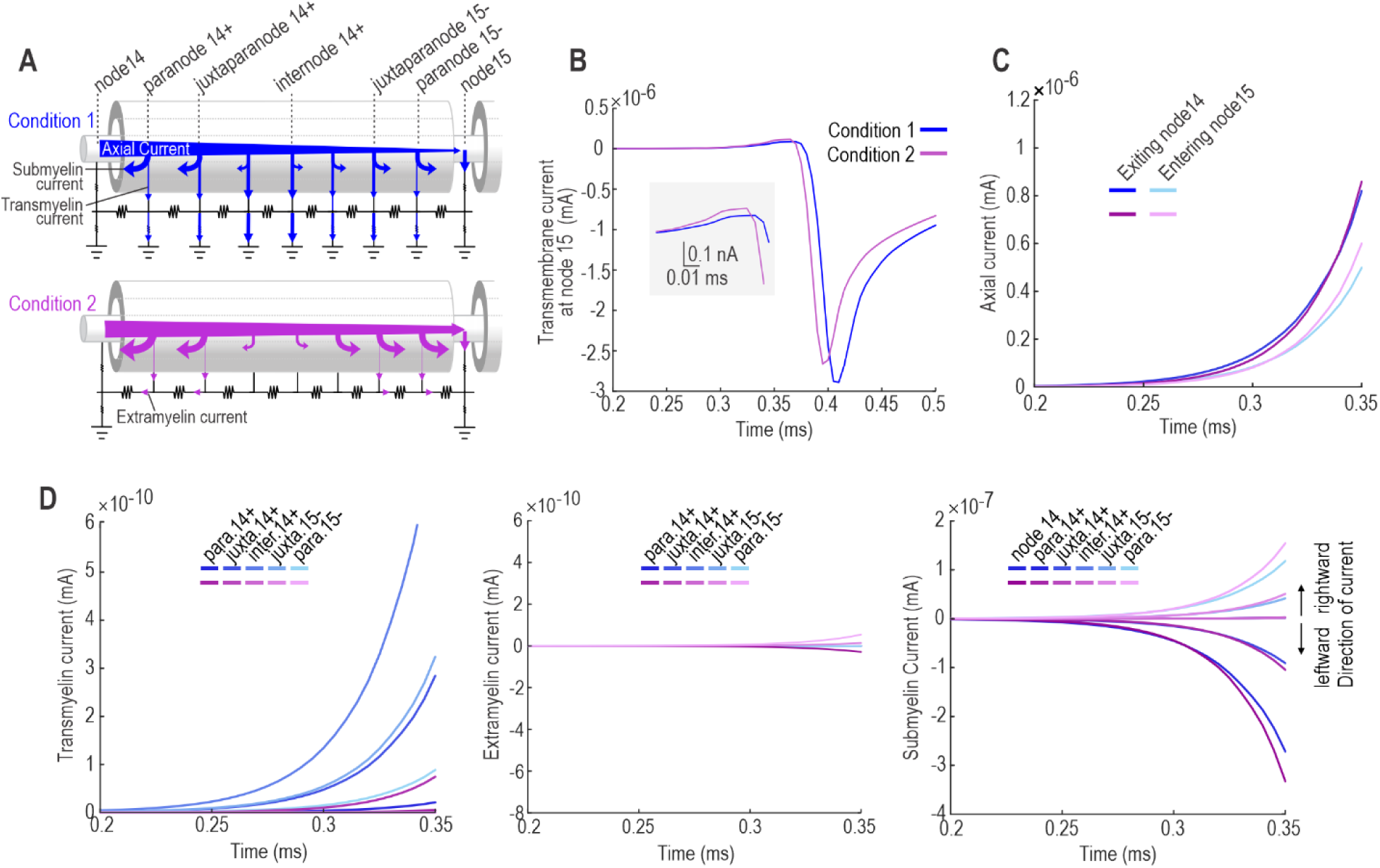
Extracellular condition 2 speeds up conduction (relative to condition 1) by shunting less axial current to the extracellular space. For panels B-D, the spike at node 14 is aligned in time for condition 1 and 2 so that any difference in spike timing at node 15 is due to effects between node 14 and 15. **A.** Cartoon illustrating axial, submyelin, transmyelin and extramyelin current in each condition. Arrow sizes approximate current amplitudes, as quantified in subsequent panels. **B.** Node 15 charges slightly faster in Condition 2, as evidenced by the larger positive phase of the transmembrane current (shaded inset), leading to earlier activation of Na channels (negative phase). **C.** The axial current leaving node 14 is comparable between the two conditions, but less axial current reaches node 15 in condition 1. This suggests that more current is lost to the extracellular space in condition 1 than in condition 2. Though the difference in axial current is small, differences in spike latency at each node accumulate to produce a large effect on conduction velocity (see Fig. 1C). **D.** Transmyelin current (left panel) was much larger in condition 1, which accounts for the difference in submyelin current between conditions 1 and 2 (right panel). Submyelin current can travel leftward or rightward depending on the longitudinal position (see cartoon in A). Note also that submyelin current is 1000x larger than transmyelin current, but 10x smaller than axial current. Extramyelin current (center panel) is negligible in both conditions; this is true even for condition 1, despite the higher transmyelin current, because current is shunted to ground.

Figure 3A compares current flow between Condition 2 (fast) and Condition 3 (slow). In Condition 3, node 15 charges slightly slower, causing Na channels to activate later (Fig. 3B). This is due to a greater attenuation of the axial current (Fig. 3C). Axial current is more attenuated in Condition 3 because more current escapes through the myelin into the extramyelin space, whose resistivity is lower than in Condition 2 (Fig. 3D **left**). The heightened transmyelin current is reminiscent of Condition 1, but unlike in Condition 1 where current is shunted straight to ground, current flows longitudinally in the extramyelin space in Condition 3 (Fig. 3D **middle**). More current flows in the submyelin space in Condition 2 due to the reduced transmyelin current (Fig. 3D **right**).

**Figure 3.**
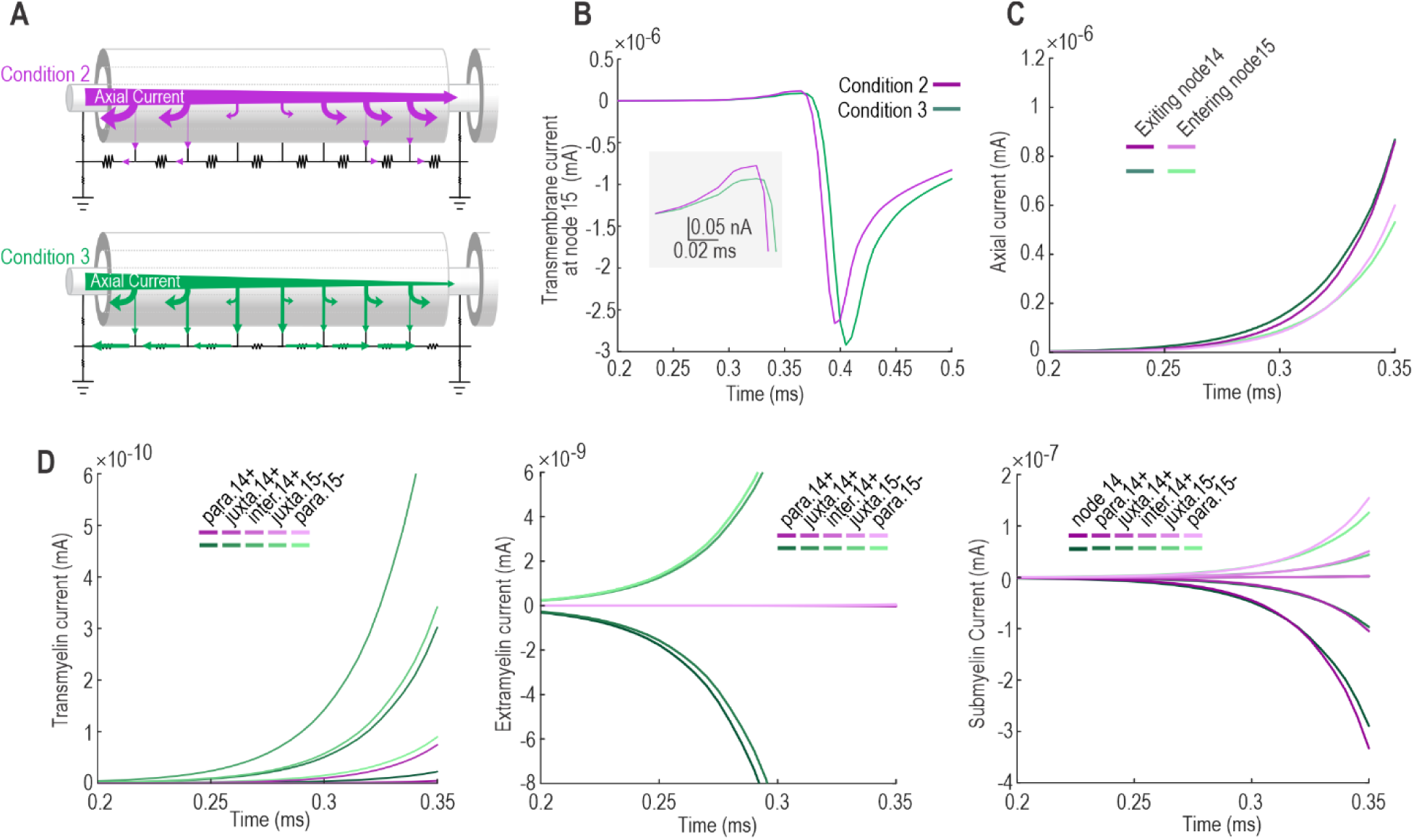
Extracellular condition 3 slows down conduction (relative to condition 1) by shunting more axial current to the extracellular space. Spikes at node 14 are temporally aligned like in Figure 2. **A.** Cartoon illustrating axial, submyelin, transmyelin and extramyelin current in each condition. **B.** Node 15 charges slightly slower in Condition 3, as evidenced by the smaller positive phase of the transmembrane current (shaded inset), leading to later activation of Na channels (negative phase) than in Condition 2. **C.** The axial current leaving node 14 is comparable between the two conditions, but less axial current reaches node 15 in condition 3. **D.** Transmyelin current (left panel) was much larger in condition 3, which accounts for the difference in submyelin current between conditions 2 and 3 (right panel). Thus far, condition 3 resembles condition 1 (see Fig. 2), but whereas transmyelin current is shunted to ground in condition 1, extramyelin current is substantial in condition 3. Extramyelin current is higher in condition 3 than in condition 2 because longitudinal resistance of the extracellular space is lower in condition 3.

According to Figures 2 and 3, the difference in conduction velocity reported in Figure 1 boils down to the amount of axial current: The more axial current reaches the next node, the faster that node charges and the earlier voltage-gated Na activate, resulting in faster saltatory conduction. Axial current is known to depend on intrinsic factors like axial resistivity and axon diameter, but even if those factors are equivalent, current flow in the extracellular space influences how much current remains in the axon and how much leaks out, some of which crosses the myelin. Transmyelin current is high if the extramyelin space is shunted straight to ground (Condition 1) or has low longitudinal resistivity (Condition 3), ultimately slowing conduction velocity. By comparison, insulation of the extramyelin space (Conditions 2) mitigates transmyelin current, leaving more current to flow axially down the axon, in which case conduction velocity is relatively fast.

### Multi-fiber model validates parameterization used in single-fiber model

In subsequent simulations, we connected the extramyelin space of each fiber to the extramyelin space of an adjacent fiber or to a fascicle boundary, thus implementing the same conditions depicted in Figure 1A but in a more anatomically realistic manner. Figure 4A illustrates the equivalent electrical circuit of the multi-fiber model including two fibers positioned closely to each other. Since the extramyelin space over internodes is disconnected from ground and the extracellular space over each node is connected to the extracellular space over the node of an adjacent fiber via a transverse resistance, this model approximates condition 2, and is henceforth referred to as Condition 2*. Conduction velocity in Condition 2* is higher than in Condition 1 due to more axial current being received by node 15, causing earlier activation of Na channels (Fig. 4B), reminiscent of results in Figures 2 and 3.

**Figure 4.**
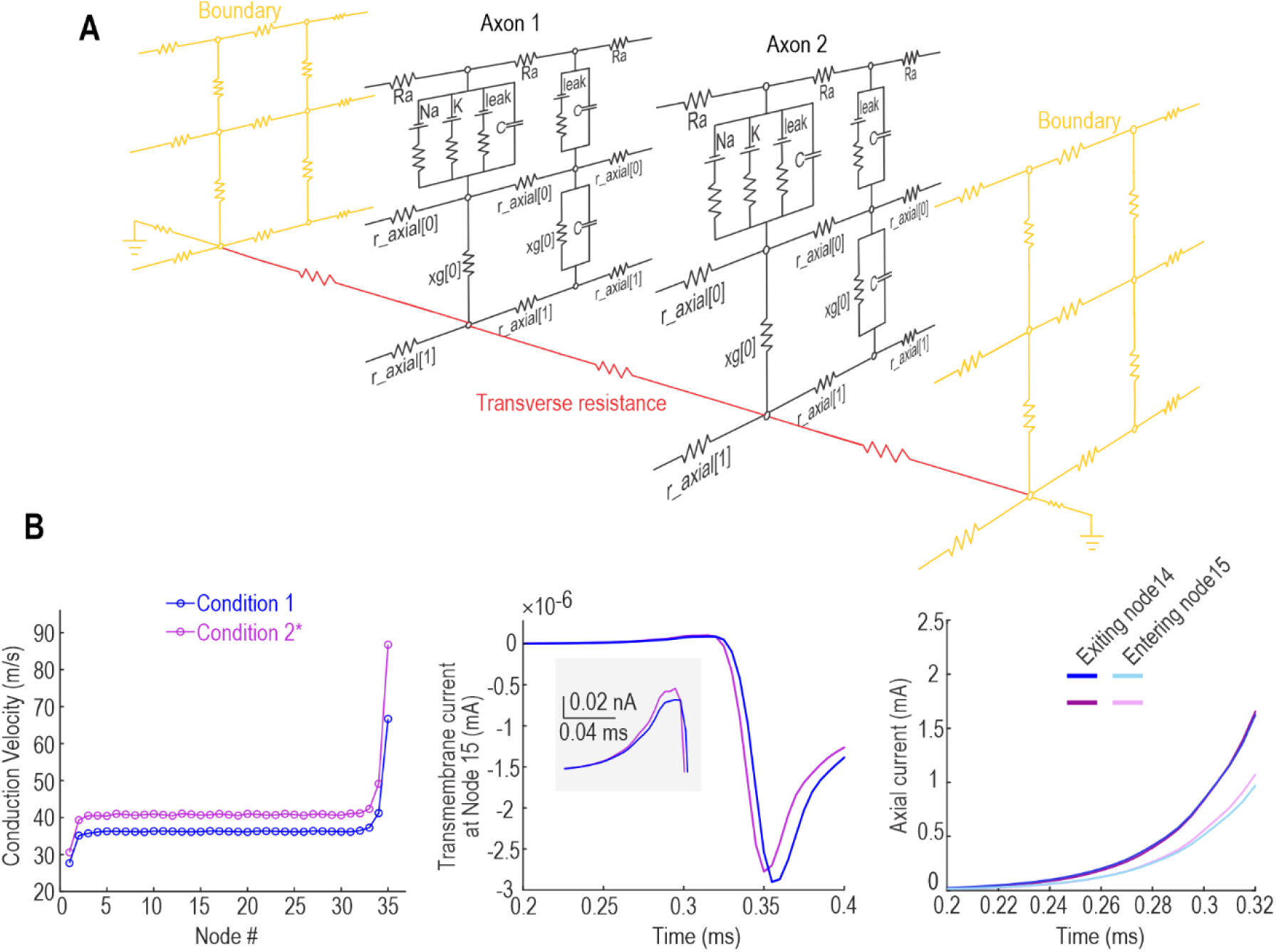
Equivalent electrical circuit of multi-fiber model. **A.** Equivalent electrical circuit for multi-fiber model. The extramyelin layers of each fiber are disconnected from ground and, instead, are connected via transverse resistances (red) to each other and to boundary cables representing the fascicle wall (gold). This is referred to as condition 2* since it is a more sophisticated variant of the single-fiber model in condition 2. **B.** Conduction velocity in Condition 2* is higher than Condition 1 (left panel) for the same reasons explained before: greater axial current reaches the next node (right panel) in condition 2, thus causing the node to charge faster and activate sodium channels sooner (center panel).

Using the multi-fiber model, we altered the anatomical properties of the extracellular space to investigate effects on conduction velocity. Two extracellular anatomical features were examined: the spacing between the fibers and the alignment of the nodes of Ranvier (Fig. 5A). Increasing the spacing between fibers or misaligning their nodes both reduced conduction velocity (Fig. 5B, C). Notably, misaligning nodes has a greater impact on conduction velocity when fibers are tightly packed because current flowing longitudinally through the extracellular space to eventually reach ground, which contributes only when nodes are misaligned, experiences more resistance when fibers are tightly packed; in other words, the relative ease of longitudinal current flow between loosely packed fibers mitigates the effect of node misalignment. Results of multi-fiber simulations are thus entirely consistent with initial single-fiber simulations but the interaction between node alignment and fiber spacing raises the possibility of other such interactions, and specifically that certain intrinsic factors may be more or less important for conduction velocity depending on extrinsic factors.

**Figure 5.**
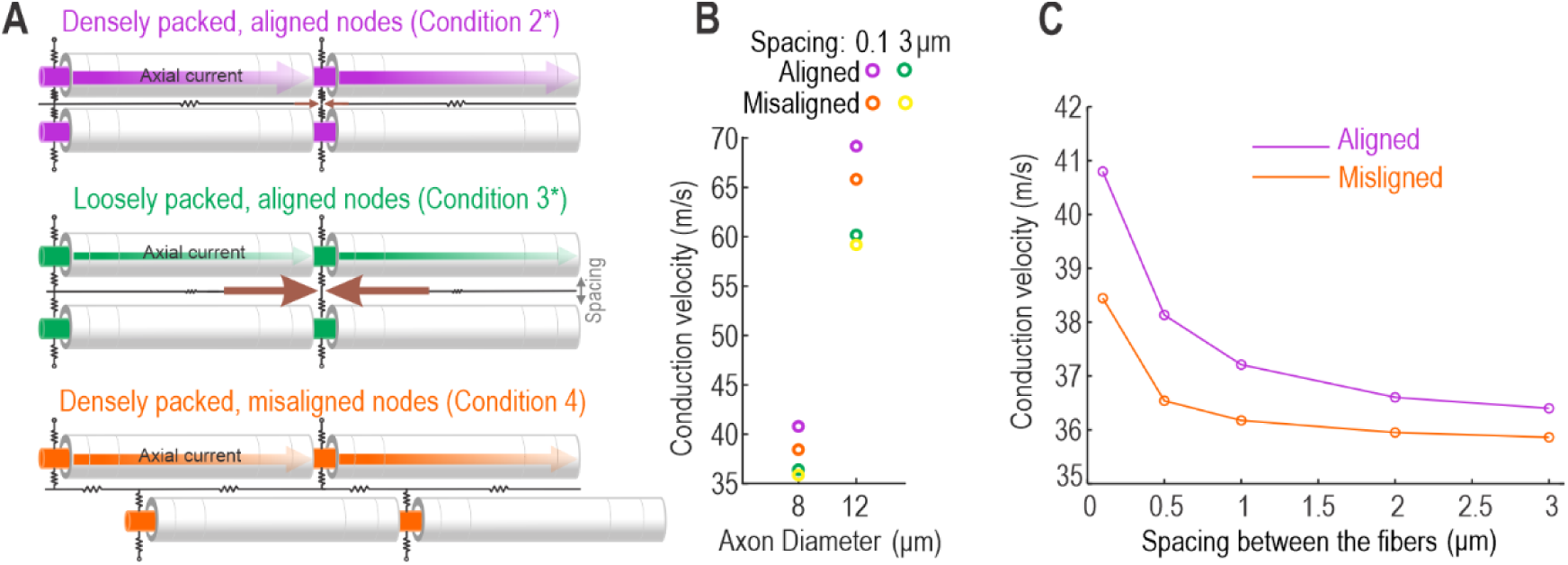
Extracellular factors interact to control conduction velocity. **A.** Cartoon illustrating different combinations of packing density and node alignment. Condition 3* represents multi-fiber variant of condition 3, and condition 4 considers misaligned nodes. **B.** Larger fibers have higher conduction velocities but, for the same axon diameter, conduction velocity is faster for densely packed axons (compare 0.1 and 3 μm spacing). Misaligning the nodes also reduces conduction velocity, but more so for tightly packed fibers (purple vs orange; see also panel C) than for loosely packed fibers (green vs yellow). **C.** Relationship between conduction velocity and fiber spacing. The gap between the two curves shows the effect of node alignment.

### Interaction between intrinsic and extrinsic properties

To test if the impact of intrinsic properties depends on extracellular conditions, effects on conduction velocity of node diameter, axon diameter, internode length, axial resistivity, number of myelin sheaths, resistivity of the periaxonal space, and resistivity of the paranode were re-tested under different extracellular conditions (Fig. 6A). Results show that conduction velocity is sensitive to axon diameter, axial resistivity, internode length, and paranode resistivity. Notably, increasing axon diameter or internode length increased conduction velocity more in Condition 2* than in Condition 1 (Fig. 6B) because axial current to the next node is especially efficient if extracellular resistance is high (Condition 2*) *and* intracellular resistance is low. Reducing axon diameter or internode length had a similar slowing effect regardless of extracellular conditions.

**Figure 6.**
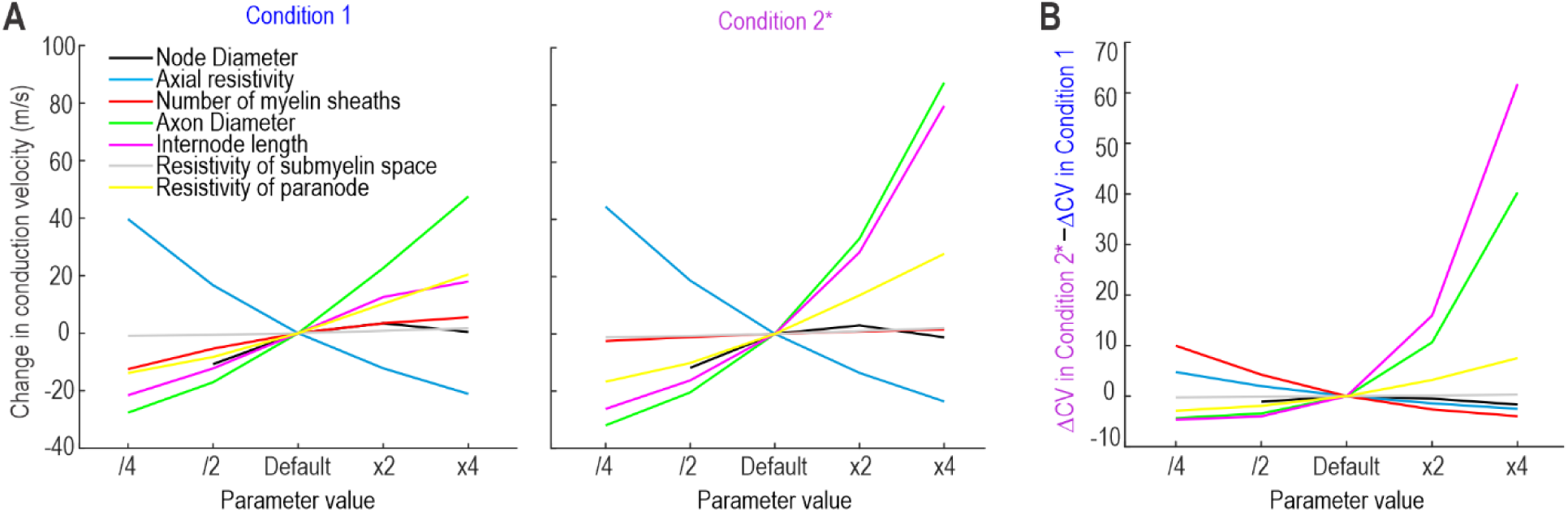
Effect of extracellular conditions on the influence of intrinsic factors on conduction velocity. **A**. Change in conduction velocity caused by increasing or decreasing a parameter of interest by 2- or 4-fold. This was repeated for condition 1 (left) and condition 2* (right). Conduction velocity is especially sensitive to axon diameter (green), axial resistivity (blue), internode length (pink), and paranode resistivity (yellow) under both conditions, but to different degrees. And whereas the number of myelin sheaths (red) affects conduction velocity in condition 1, it has little effect in condition 2*. **B.** The interaction between parameter variations and extracellular conditions is summarized by plotting the difference between conditions (i.e. the difference in differences). Whereas axial resistivity has a similar large effect in both conditions (resulting a in a flat difference of difference curve), increasing axon diameter and internode length has a larger effect in condition 2* (resulting in steep difference of difference curves).

On the other hand, the number of myelin sheaths impacted conduction velocity in condition 1 but has remarkably little effect in condition 2*, as documented more thoroughly in **Figure 7**. Reducing the number of myelin sheaths caused a dramatic slowing and eventual failure of conduction in Condition 1 but not in Condition 2* (**Fig. 7A,B**). This occurs because, in condition 2*, the intact myelin of adjacent fibers provides a “backup” layer of insulation that mitigates the flow of extracellular current to ground. The energy efficiency of spike propagation is also differentially sensitive to the number of myelin sheaths contingent on extracellular conditions (**Fig. 7C**). Energy efficiency was calculated (**Hu et al., 2018**):

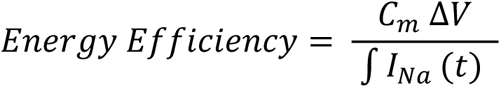

where *C*_*m*_ is specific membrane capacitance of the node, Δ*V* is the maximum voltage deflection of the spike, and *I*_*Na*_ is the sodium current. According to this metric, an efficient axon uses the minimal sodium current required to drive the transmembrane voltage change.

**Figure 7.**
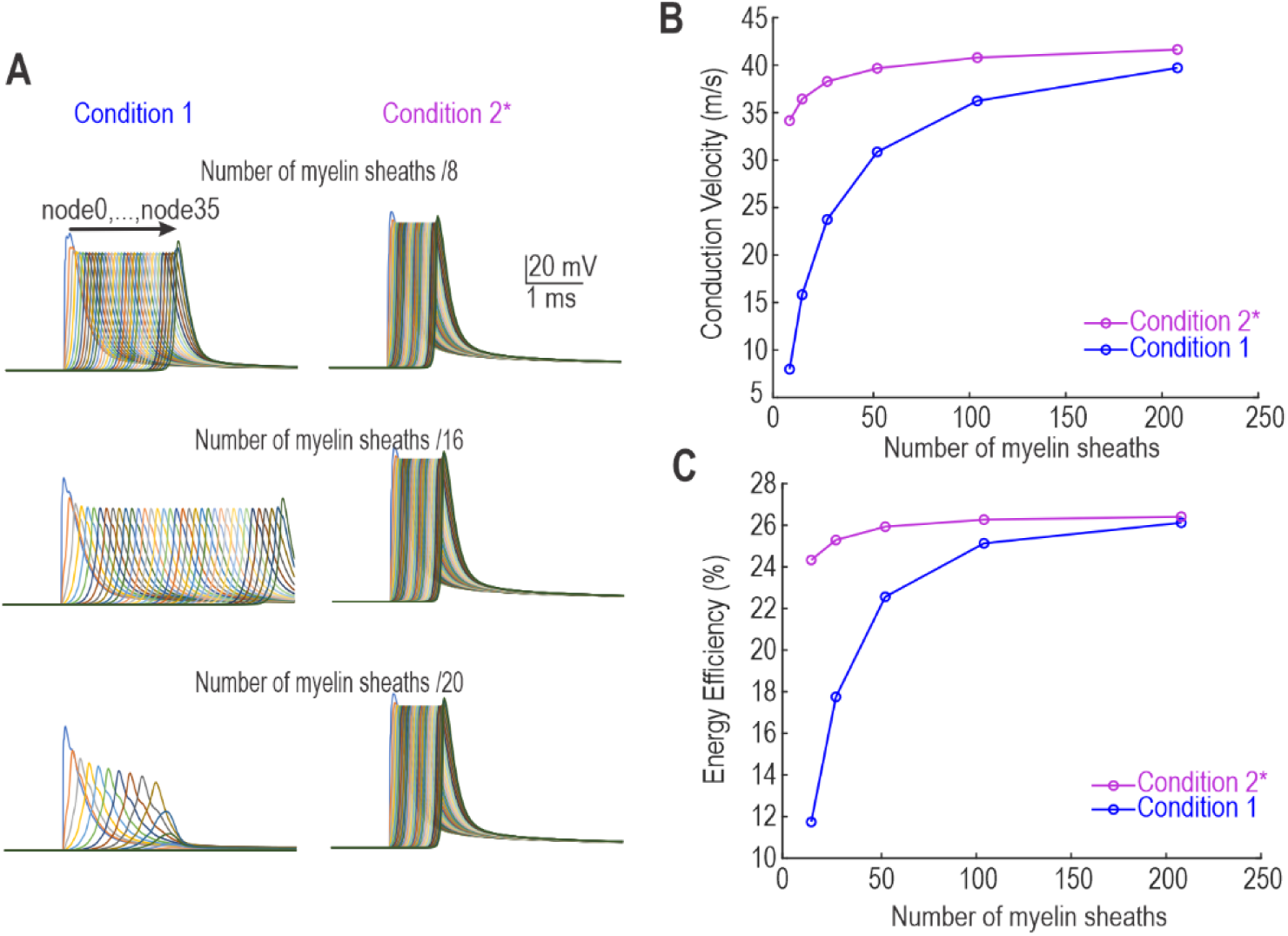
Effect of myelin thickness on conduction velocity, reliability and energy efficiency depends on extracellular conditions. **A.** Spikes from all nodes overlaid to show propagation over time for different numbers of myelin sheaths. The default number of sheaths is 104. In condition 1 (left), decreasing the number of sheaths dramatically slowed conduction and eventually caused propagation to fail, whereas the same change had relatively little effect in condition 2* (right). **B.** Conduction velocity plotted as a function of myelin thickness. **C.** Energy efficiency of spike generation also depends jointly on the number of myelin sheaths and extracellular conditions.

Overall, these results demonstrate that the importance of any one parameter on axon function depends on other parameters, and that this interdependence can be understood in terms of intracellular and extracellular currents.

### Ephaptic effects also depend on extracellular conditions

In Condition 2*, we calculated the extramyelin current flowing from one fiber that was propagating a spike to a nearby fiber that was quiescent. The extracellular current caused by a spike in the first fiber produced a small subthreshold voltage change in the nearby fiber (Fig. 8). The current waveform flowing from the active fiber to the quiescent fiber is biphasic, which results in the latter experiencing a biphasic extracellular voltage change, which translates to a biphasic transmembrane voltage change. Specifically, initial subthreshold depolarization in the active fiber (caused by axial current arriving from the upstream node) causes a current through the transverse resistance that effectively hyperpolarizes the quiescent fiber by causing its extracellular space to become more positively charged. Subsequent suprathreshold depolarization in the active fiber (caused by inward current through activated Na channels in the node) causes the opposite transverse current flow, leading to depolarization of the quiescent axon. Notably, the extracellular potential corresponds to the spike waveform that can be recorded extracellularly; this particular waveform is not triphasic because the afterhyperpolarization is absent from transmembrane voltage waveform since axonal spike repolarization is mediated primarily by Na channel inactivation rather than by delayed rectifier K channel activation. These results are not novel but, instead, confirm that our model works as expected.

**Figure 8.**
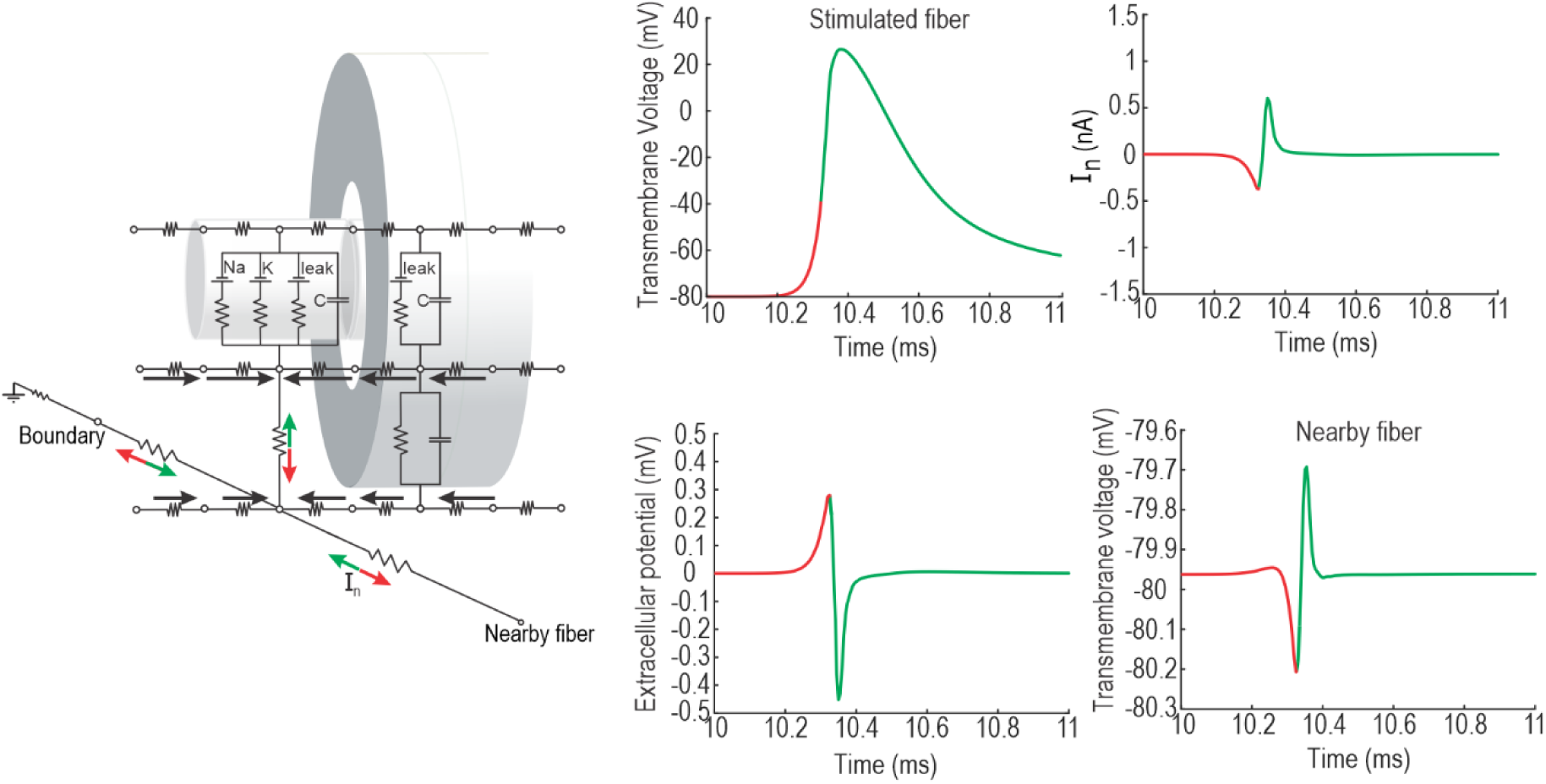
A spike in one axon causes subthreshold voltage changes in a nearby axon. Equivalent circuit diagram highlights extracellular current moving away (red arrows) or toward (green arrows) the stimulated fiber that propagates an action potential. The same color convention is used in graphs to distinguish different phases of the response. Black arrows highlight direction of longitudinal current in submyelin and extramyelin spaces. As the node charges due to incoming axial current (red phase of top left graph), the extracellular voltage increases (bottom left graph), resulting in extracellular current flowing away from the activated node (top right graph), causing hyperpolarization in a nearby node (bottom right graph). As sodium channels activate (green phase), the extracellular voltage and extracellular current flow reverse direction, causing depolarization in the nearby fiber.

To explore how extracellular factors affect ephaptic interactions with nearby fibers, we increased the number of simulated fibers to 20. All fibers had the same intrinsic properties and were connected to each other and to the boundary via transverse resistances with different values **(Fig. 9A).** A spike was evoked at the end of one fiber positioned in the middle of the fascicle, and the peak-to-peak transmembrane voltage changes in all other (quiescent) fibers were calculated; this was repeated with the nodes aligned or misaligned (by randomly shifting fibers longitudinally). As shown in **Fig. 9B**, the ephaptic voltage change in quiescent fibers was not sensitive to the distance from the active fiber, but was sensitive to node alignment; specifically, node misalignment diminished ephaptic effects. In **Fig. 9C**, a spike was evoked synchronously in 19 fibers and the effect on one quiescent fiber was examined. As expected, the cumulative ephaptic effect of 19 active fibers produced a transmembrane voltage change ∼10x larger than that caused by a spike evoked in one fiber, but node misalignment still diminished the effect.

**Figure 9.**
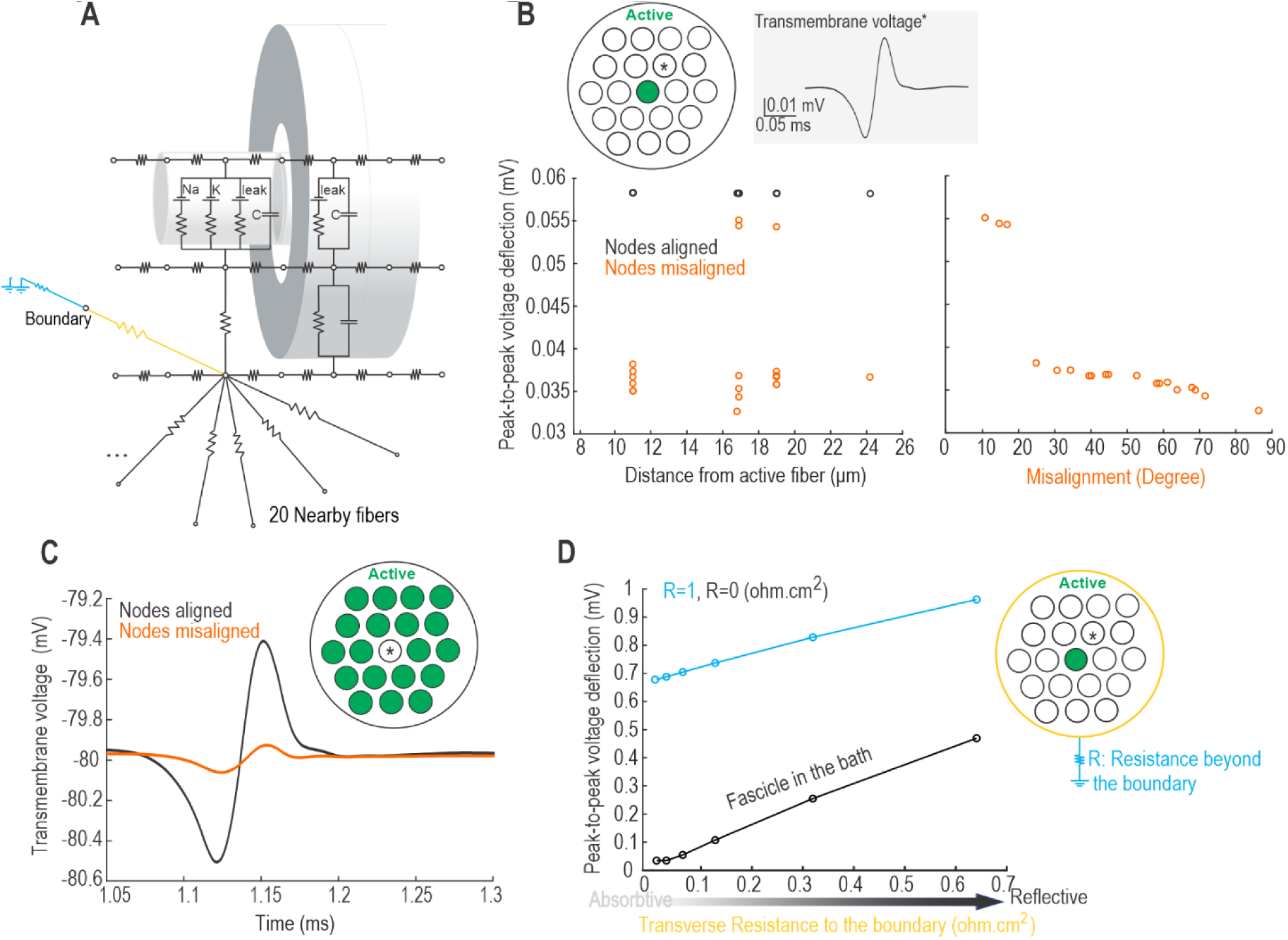
Ephaptic interactions are influenced by extracellular conditions. **A.** Equivalent circuit diagram showing increased lateral connections to 20 nearby fibers. All fibers have the same intrinsic properties. **B**. Effect of a spike in one fiber (green) on nearby fibers. Grey inset shows sample voltage deflection in nearby fiber (*). The simulated fascicle included 20 fibers spaced 3 μm edge-to-edge, on average, from their nearest neighbors. The ephaptic voltage deflection in nearby fibers is independent of distance from the activated fiber (left panel) regardless of node alignment. Increased variability for misaligned nodes (orange) is accounted for by degree of misalignment (right panel). **C**. Effect of a synchronous spike in 19 fibers (green) on a nearby fiber (*). The cumulative ephaptic effect of 19 activated fibers is much larger than the unitary ephatic effect in panel B, but is still diminished by node misalignment. **D**. Effect of resistance to the perineurium (gold) and beyond (blue). Increasing or decreasing transverse resistances render the boundary more reflective or absorptive, respectively. Voltage deflection in nearby fibers is increased by a more reflective boundary condition. Moreover, if the extraperineurial space is simulated as if in a bath (R=0, black curve), ephaptic effects are smaller than if R is larger (blue).

To test the effects of the boundary condition surrounding the fibers, the transverse resistances between the axons and the boundary cables were changed (Fig. 9D). Large and small transverse resistances render the boundary relatively more reflective or absorptive, respectively. A more reflective (insulating) boundary condition increased ephaptic effects by delivering more charge to adjacent fibers. Likewise, increasing the resistivity of the extracellular space outside the fascicle increased ephaptic effects (compare blue and black curves). These results demonstrate the importance of boundary conditions, and highlight the caveats of simulating single fibers in isolation.

## DISCUSSION

Results of the current study reveal that, alongside factors intrinsic to the fiber such as axon diameter and myelin thickness, extracellular conditions also affect saltatory spike propagation by influencing where current flows. Specifically, using computational simulations, we measured current passing through various intracellular and extracellular pathways, which is infeasible to fully assess experimentally. We found that conduction velocity of a myelinated axon is slower when extracellular current is shunted to ground (i.e. boundary conditions are absorptive) whereas conduction is faster if extracellular conditions are more resistive (Fig. 1). This is explained by effects on axial current: the axial current flowing from one node to the next is diminished by loss of current to the extracellular space, which ultimately depends on the extracellular boundary conditions. The more current is lost from the intracellular space, the slower the next node charges, thus delaying spike regeneration (Figs. 2 and 3). The resistance encountered by extracellular current flow depends on factors such as fiber packing density, where denser packing conveys greater resistivity (Fig. 5). But one must of course also consider positioning of a fiber within a fascicle, since fibers positioned at the edge of a densely packed fascicle do not benefit from the same total extracellular resistance as fibers squeezed in the middle. These results have several other consequences, as discussed below.

Our findings highlight the importance of considering extracellular conditions when examining the impact of intrinsic properties on conduction velocity, as the former can diminish or amplify effects of the latter (see Fig. 6). For example, in a less absorptive extracellular condition, the impact of demyelination on spike propagation is considerably reduced (Fig 7). Under pathological demyelinating conditions, several fibers are likely to experience demyelination simultaneously, in which case any one demyelinated fiber cannot rely on the myelin of neighboring fibers. But under normal conditions, unmyelinated fibers are positioned amongst myelinated fibers and may benefit from the electrical insulation this affords. Indeed, unmyelinated fibers associate with Schwann cells, forming Remak bundles (Harty & Monk, 2017), but resistance to extracellular current flow beyond this immediate ensheathment is liable to increase the speed and energy efficiency of continuous (non-saltatory) spike propagation, by maximizing axial current. While previous modeling studies have explored the effects of myelin thickness and demyelination on spike propagation (Lasiene et al., 2008; Powers et al., 2012; Scurfield & Latimer, 2018), our study highlights the importance of considering extracellular conditions.

Most computational studies have investigated ephaptic coupling by modeling identical aligned fibers to reduce computational cost (Binczak et al., 2001; Reutskiy et al., 2003). We show that spatial relationships between nodes affect ephaptic coupling; specifically, node misalignment decreases ephaptic coupling among axons (Fig 9). Our finding is consistent with those of Capllonch-Juan et al. (2020), who found that fiber heterogeneity and node misalignment diminish ephaptic effects. Furthermore, we emphasize the importance of boundary conditions on ephaptic effects, where a more absorptive boundary condition reduces ephaptic effects (Fig. 9). This is consistent with early experiments indicating that when axons are exposed to a solution with lower salinity, the conductivity of extracellular fluid decreases, causing increased ephaptic coupling (Katz & Schmitt, 1940); placing the axons in oil has a similar effect (Jefferys, 1995; Ramon & Moore, 1978).

It should be intuitive that if more extracellular current is promptly lost to ground, then less current is available to interact with adjacent fibers. The same logic applies to the amplitude of extracellularly recorded spikes, depending of course on the position of the recording electrode relative to the active fiber and to various sources of extracellular resistance. Interestingly, ephaptic interactions tend, on average, to reduce conduction velocity (Capllonch-Juan & Sepulveda, 2020; Schmidt & Knösche, 2019); we have shown that more resistive boundary conditions increase conduction velocity but, concurrently, increase ephaptic coupling, which could indirectly reduce conduction velocity if spikes synchronize (Schmidt et al., 2021). Separate from affecting spike propagation in axons, ephaptic interactions can influence spike initiation (in the soma or axon initial segment) and thereby influence the likelihood or timing of spike initiation during network oscillations or otherwise synchronized synaptic input (Anastassiou et al., 2011; Fröhlich & McCormick, 2010; Goldwyn & Rinzel, 2016; Greg Stacey et al., 2015). In both scenarios, sizeable effects normally rely on coordinated activity across multiple neurons (Schmidt et al., 2021; Vigmond et al., 1997).

The importance of the extracellular space and the diffusion barrier created by Schwann cells has long been recognized as an explanation for the accumulation of extracellular potassium during sustained neuronal activity (Frankenhaeuser & Hodgkin, 1956; Taylor et al., 1980) since potassium ions are released into the extracellular space during spike repolarization. Elevated extracellular potassium levels lead to changes in the resting membrane potential and increased neuronal excitability (Contreras et al., 2021). Under normal conditions, extracellular potassium is efficiently cleared away by various mechanisms, including diffusion through the extracellular space (Gardner-Medwin, 1983; Taylor et al., 1980). The diffusion barrier formed by Schwann cells, along with other factors such as the presence of blood vessels, influences the properties of the extracellular space, thereby affecting the clearance of extracellular potassium ions. Effects of restricted ion movement can accumulate over long timescales, depending on activity levels, leading to obvious consequences for axon function; the same processes occur on faster timescales, with subtler but not necessarily negligible effects.

Our results are also notable for efforts to infer axon properties by fitting models to experimental data. This is increasingly feasible thanks to increasing computing power and improved optimization methods (e.g. Van Geit et al., 2016), but there is usually no unique solution, rendering it very difficult to identify which particular solution is used by a given axon. Identifying the range of possible parameters that could yield the observed data may suffice for some purposes but will not, however, enable accurate predictions of how that axon will respond to a particular parameter change such as thinning of the myelin sheath. Considering many constraints (e.g. measuring conduction velocity *and* fiber diameter) can mitigate this problem, but the fact remains that robust biological systems tend to be degenerate, meaning they have multiple viable solutions available to them so that compensation is possible (Yang et al., 2022; Yang & Prescott, 2023). Established co-variations between certain factors – like the g-ratio, which relates axon diameter to total (outside) fiber diameter and implies a relationship between axon diameter and myelin thickness (Rushton, 1951) – restricts the solution space. But the solution space is expanded by our observation that extracellular conditions (which might themselves vary) also influence conduction velocity. It is notable that certain interactions are multiplicative rather than additive; for example, the effect of myelin thickness on conduction velocity is scaled by extracellular conditions (Fig. 6).

In conclusion, these results of this study highlight the relatively unappreciated influence of extracellular conditions on saltatory spike propagation in myelinated axons. The speeding up or slowing down of propagation is explained by changes in the amount of axial current, which depends on how much current is lost to the extracellular space, depending on extracellular conditions. The flow of extracellular current also affects ephaptic interactions. The effects of extracellular current flow are usually modest but may, under certain circumstances, become important and should, therefore, merit attention.

## Conflict of interest

None

## Acknowledgments

This work was funded by the Canadian Institutes of Health Research (CIHR) Foundation Grant 167276 to SAP.

## Notes

### Competing Interest Statement

The authors have declared no competing interest.

